# Direct Bioconjugations of Diverse Payloads using an Extraordinarily Donor-Promiscuous Flavin Transferase

**DOI:** 10.64898/2026.10.31.763105

**Authors:** Wahyu S. Widodo, Xiaoman Fan, Ruben R. Kamphuis, Karla L.H. Feijs-Žaja, Alexander Belyy, Maximilian J.L.J Fürst

## Abstract

Site-specific protein conjugation is a widely used tool in chemical biology research and is the foundation of many biotechnological and biomedical applications. A key enabling technology are repurposed post-translational protein modifying enzymes, where variants with broad donor and acceptor substrate promiscuity were developed into general-purpose bioconjugation platforms. However, direct enzymatic attachment of bulky or structurally diverse payloads is often limited by low reaction efficiencies and difficulties in preparing suitable donors. Consequently, conjugation typically relies on two-step strategies in which an enzyme first installs a bioorthogonal handle on the target protein, followed by chemical attachment of the payload. These enzyme-first, chemistry-second approaches limit chemistry to bioorthogonal reactions that leave a molecular footprint. Here, we introduce direct ADDing, an enzymatic bioconjugation platform based on our discovery of the exceptional donor promiscuity of the flavin transferase ApbE. We show that ApbE accepts a remarkably broad range of donor substrates provided they carry an adenosine diphosphate (ADP) anchor. The prevalence of ADP-containing compounds in nature provides access to diverse building blocks and an extensive biochemical toolbox for donor generation. Exploiting these resources, we demonstrate multiple strategies for preparing compatible donors and achieve direct, site-specific enzymatic conjugation of diverse small molecules, peptides, proteins, and synthetic polymers.

## Introduction

For decades, chemical biologists have employed bioconjugation to equip biomolecules with functions beyond their natural repertoire. A broad range of chemical and biochemical methods now enables the precise installation of functional groups on proteins of interest.^1^ Applications range from research tools, such as fluorescently labeled proteins for microscopy or protein-DNA conjugates for single-molecule studies, to biomedicines such as antibody-drug conjugates (ADCs).^2–5^

Although chemical modification of addressable protein side chains, particularly cysteines and lysines, is widely established, these approaches face limitations including undesired side reactions, heterogeneous modification, and incompatibility with complex biological samples.^1^ These challenges have driven the development of site-specific bioconjugation methods, many of which employ enzymes repurposed from natural post-translational modification (PTM) pathways. Among the most widely adopted are those based on Sortase A,^6,7^ phosphopantetheinyl transferase (PPTase),^8–10^ lipoic acid ligase,^11–13^ microbial transglutaminase,^14,15^ and the formylglycine-generating enzyme.^16^ These systems generally recognize defined sequence motifs or structural features that can be genetically introduced into target proteins, enabling selective attachment of functional molecules (payloads). Choosing among them requires balancing multiple criteria: some applications impose specific requirements on linker chemistry, whereas others prioritize short acceptor tags, experimental simplicity, or high conjugation yields. No single method satisfies all requirements simultaneously.^17^

A major limitation of enzymatic bioconjugation is that the intended payload often cannot be attached directly. Direct conjugation faces two principal obstacles: preparing payloads bearing the molecular recognition elements required by the enzyme, and ensuring that the enzyme accepts the resulting donor substrates. These recognition elements, or molecular anchors, are dictated by the enzymes’ natural functions and frequently comprise complex biomolecular structures that are challenging to introduce synthetically. Moreover, although bioconjugation enzymes are generally selected for their ability to accept non-native substrates, their tolerance to payload variation is not unlimited.^18,19^ Consequently, many conjugation strategies rely on two steps: enzymatic installation of a small bioorthogonal handle, followed by chemical attachment of the desired payload. However, this requires chemistry to be performed on the modified protein, potentially limiting compatibility with complex biological samples. Furthermore, the resulting linkage often introduces “scars” such as triazole rings, which may interfere with the intended application.

Overcoming both synthetic and enzymatic constraints would therefore expand the scope of direct bioconjugation. Its potential is illustrated by applications in which donor preparation is relatively straightforward. For example, the modularity of phosphoramidite synthesis facilitates the incorporation of conjugation anchors into oligonucleotides, enabling direct preparation of protein-oligonucleotide conjugates.^20^ Although many enzymatic systems struggle with such bulky substrates, notable exceptions include Sortase A and the PPTase Sfp. Both tolerate substantial structural variation beyond their respective recognition elements, enabling attachment of extremely large payloads.^21,22^ Nevertheless, preparing compatible donor substrates remains synthetically demanding and costly, particularly for complex molecules. Although protein engineering offers opportunities to expand substrate tolerance,^12,23^ this process is laborious and often yields variants optimized for individual modifications.^24–27^

Thus, a need remains a need for a site-specific bioconjugation system that combines facile donor preparation with exceptional tolerance to payload properties, particularly size. Here, we demonstrate that the flavin transferase ApbE from *Vibrio cholerae* meets these requirements. Previous work established ApbE as a robust platform for site-specific flavinylation through recognition of a short, seven-residue acceptor motif that can be introduced into diverse target proteins. These studies also revealed tolerance to modifications of the flavin scaffold,^28,29^ which we subsequently exploited to enzymatically install click-chemistry handles for two-step conjugation.^30^ In the present work, we investigate a reverse strategy in which payload functionalization precedes enzymatic attachment, enabling direct bioconjugation (Figure 1).We systematically varied the molecular components of ApbE’s natural donor, flavin adenine dinucleotide (FAD), to explore the limits of its substrate tolerance and suitability for direct conjugation. We found that ApbE exhibits exceptional promiscuity, accommodating a broad range of payloads presented as adenine dinucleotide derivatives (ADDs). Taking advantage of the abundance of ADP-containing compounds in nature, their commercial availability, and biochemical methods for generating and modifying them, we establish ADD-tagging as a versatile platform for direct protein conjugation, accommodating payloads ranging from small molecules and peptides to proteins and solid supports.

**Figure 1.**
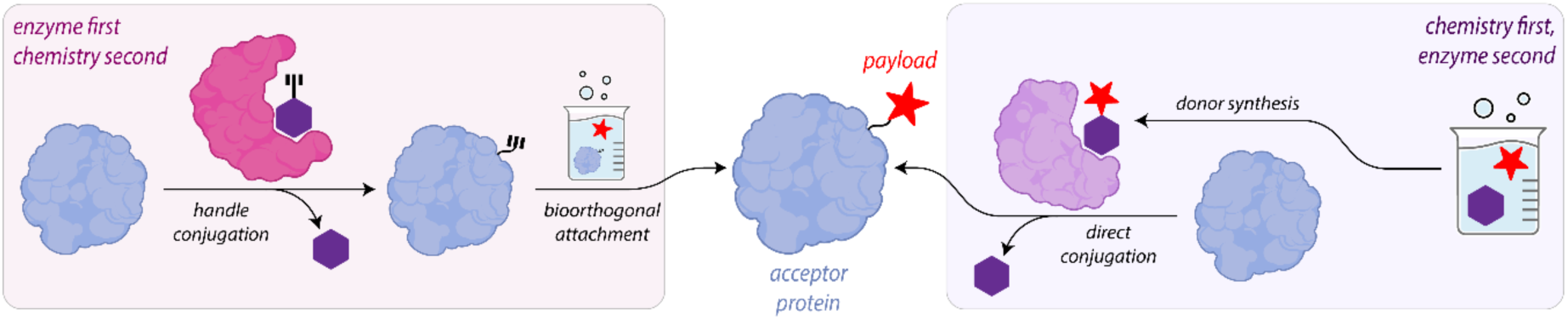
Schematic comparison of two-step (enzyme-first, chemistry-second) and direct (chemistry-first, enzyme-second) bioconjugation strategies. For challenging and bulky substrates, many approaches including our previous click ADDing,^30^ use the two-step approach, while the present study show that due to an exceptional promiscuity, the flavin transferase ApbE is also suitable for the alternative one-step approach.

## Results

### ADP Moiety Recognition Underlies ApbE Promiscuity

Previous work by us and others^29–31^ demonstrated ApbE activity with a handful of closely related donors, including FAD analogues bearing modified isoalloxazine rings (5-deaza- and roseoflavin derivatives), as well as three ADP-ribose (ADPR) derivatives: NADH, 1-O-(pent-4-ynyl)-ADPR, and 1-O-(3-azidopropyl)-ADPR (Table 1 and S1). These donors all share an ADP moiety linked via a ribityl or ribosyl to an isoalloxazine ring, nicotinamide, or a short aliphatic chain, respectively. In each case, the resulting modification is predictable: in line with its pyrophosphate-cleaving activity, ApbE links a reactive threonine or serine residue in the acceptor motif to the adenosine-distal phosphate, forming a phosphodiester bond. Although enzyme tolerance toward modest variations in a natural substrate’s molecular scaffold is not uncommon, particularly when the modifications reduce molecular size, we found it remarkable that ApbE is apparently indifferent to the structural disparity between acyclic ribityl and cyclic ribosyl linkers in close proximity to the reactive atoms.

**Table 1.**
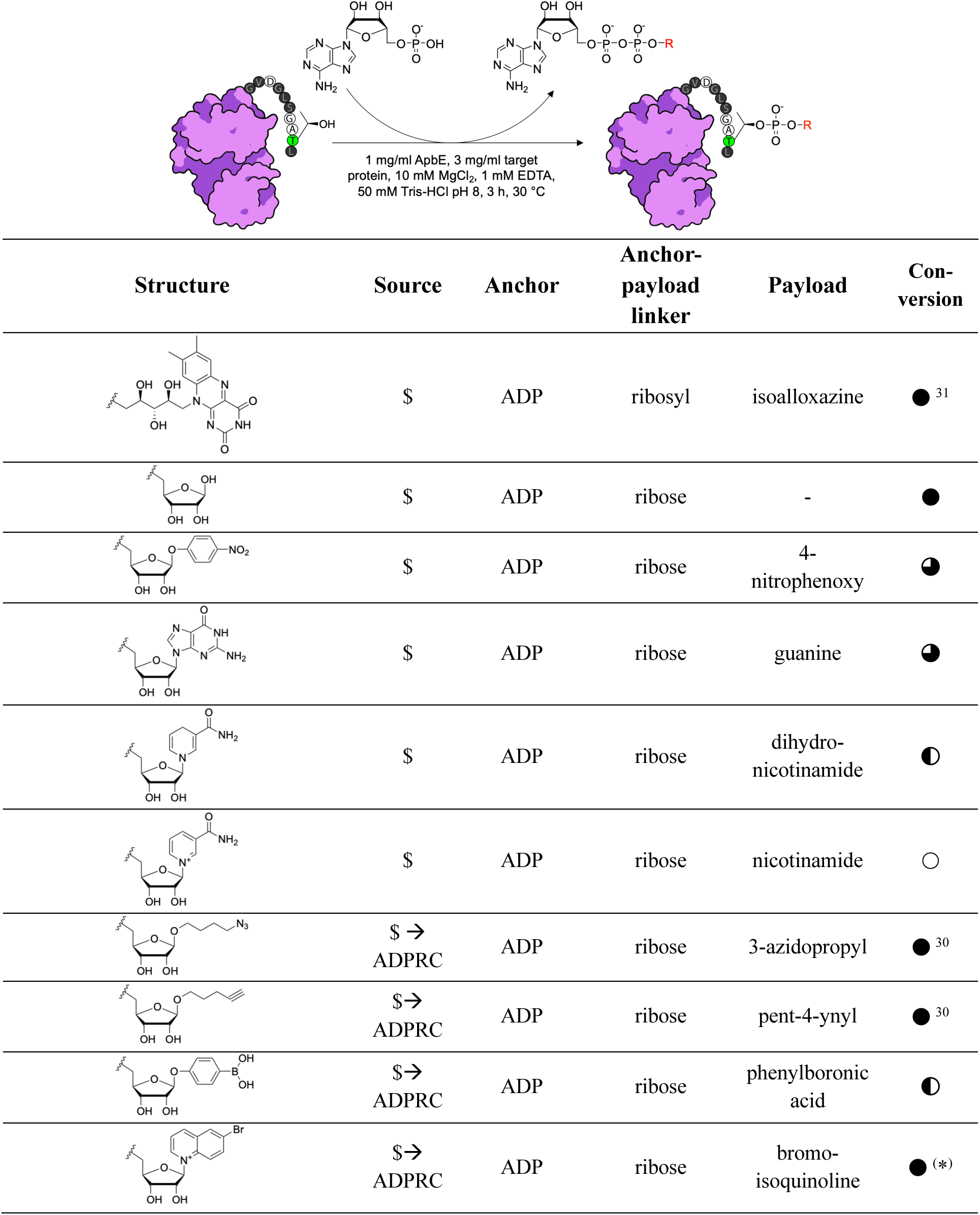
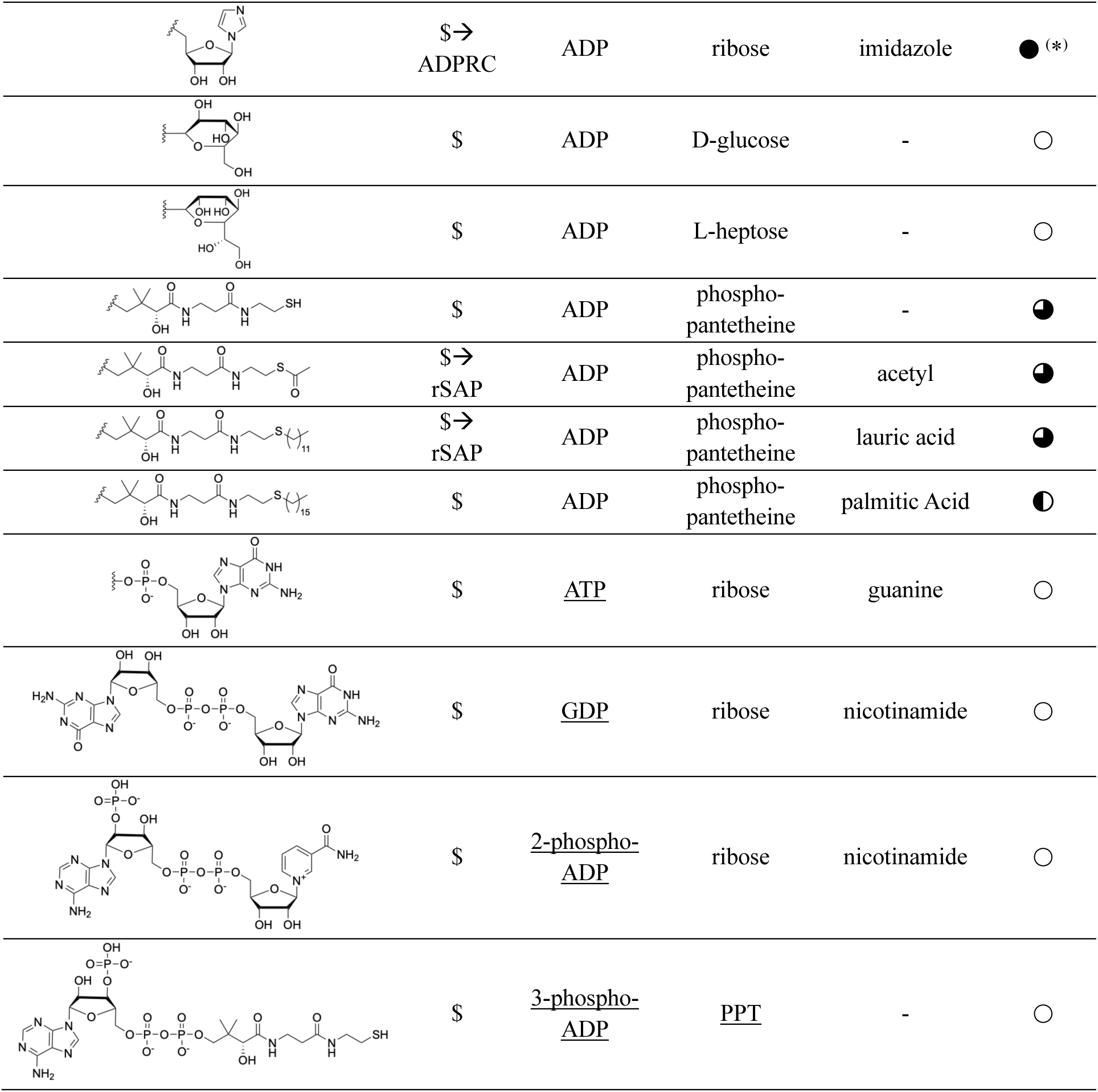
Substrate specificity of ApbE. ApbE-catalyzed protein modifications using small molecules with structural variations in anchor or payload scaffold as donors, where reactions were classified as full (⬤), > 50% (◕), ∼ 50% (◐), <50% (◔) or no conversion (◯) based on deconvoluted ESI-MS data (Table S1). Commercially obtained molecules are indicated with “$”. Deviating from the represented standard protocol, substrates made by ADPRC were reacted with ApbE in one-pot, indicated by (*). A squiggly line in the structure indicates the standard ADP anchor.

| Structure | Source | Anchor | Anchor-payload linker | Payload | Con-version |
|---|---|---|---|---|---|
|  | \$ | ADP | ribosyl | isoalloxazine | ● <sup>31</sup> |
|  | \$ | ADP | ribose | - | ● |
|  | \$ | ADP | ribose | 4-nitrophenoxy | ◕ |
|  | \$ | ADP | ribose | guanine | ◕ |
|  | \$ | ADP | ribose | dihydro-nicotinamide | ◐ |
|  | \$ | ADP | ribose | nicotinamide | ○ |
|  | \$ → ADPRC | ADP | ribose | 3-azidopropyl | ● <sup>30</sup> |
|  | \$→ ADPRC | ADP | ribose | pent-4-ynyl | ● <sup>30</sup> |
|  | \$→ ADPRC | ADP | ribose | phenylboronic acid | ◐ |
|  | \$→ ADPRC | ADP | ribose | bromo-isoquinoline | ● <sup>(*)</sup> |
|  | \$→ ADPRC | ADP | ribose | imidazole | ● <sup>(*)</sup> |
|  | \$ | ADP | D-glucose | - | ○ |
|  | \$ | ADP | L-heptose | - | ○ |
|  | \$ | ADP | phospho-pantetheine | - | ◕ |
|  | \$→ rSAP | ADP | phospho-pantetheine | acetyl | ◕ |
|  | \$→ rSAP | ADP | phospho-pantetheine | lauric acid | ◕ |
|  | \$ | ADP | phospho-pantetheine | palmitic Acid | ◐ |
|  | \$ | ATP | ribose | guanine | ○ |
|  | \$ | GDP | ribose | nicotinamide | ○ |
|  | \$ | 2-phospho-ADP | ribose | nicotinamide | ○ |
|  | \$ | 3-phospho-ADP | PPT | - | ○ |

To systematically investigate the molecular requirements for ApbE-mediated donor acceptance, we sought to probe the importance of each sub-moiety and define the enzyme’s limits. We thus evaluated a panel of commercial molecules under reaction conditions we deliberately kept suboptimal, to prevent full conversions from discerning subtler differences. For these tests, we used the native acceptor protein NqrC or maltose binding protein (MBP) harboring the ADD peptide tag on the C- or N-terminus, and semi-quantitatively analyzed modifications by electrospray ionization mass spectrometry (ESI-MS). Given the four structural elements that comprise the native ApbE substrate, FAD—adenosine, a central diphosphate, a ribityl linker, and an isoalloxazine ring—we tested analogues in which these features varied to determine their significance.

We first tested modifications to the non-transferred ADP anchor. Replacing adenine with guanine, as in the symmetric Gp₂G molecule, led to no detectable product; neither did substrates with additional phosphate groups on the anchor ribose, NADPH and Coenzyme A (CoA), nor substrates with a tri-instead of a diphosphate bridge, as in Ap_3_G (Table 1, Table S1). As these negatives could also have been caused by the non-native payload rather than the altered anchor scaffold, we simultaneously tested the same chemistries with a fixed ADP anchor to probe transfer group promiscuity. While it was already known that NADH was accepted by ApbE, we found that Ap_2_G as well as dephospho-CoA (dCoA) were in fact very efficiently used by the enzyme, too. Side-by-side, these results indicate a strict requirement of ApbE for the ADP anchor, while tolerating significantly different payload chemistries. Given CoA’s importance as a central metabolic acyl group-carrier, various substituted thioester variants are commercially available and provided us with a convenient test of larger payloads. While only available as phosphorylated CoA derivatives, we could conveniently make the dCoA variants using commercial shrimp alkaline phosphatase (rSAP), and found that ApbE also installs acetyl, lauryl, and palmitoyl PPT. Given the pantetheine arm’s strikingly different architecture compared to the FAD/NADH ribityl and ribosyl, these results showed a broad tolerance of ApbE both toward the donor chemistry directly adjacent to the attachment point, as well as to bulkier payloads. Interestingly, we did encounter some limitations in ApbE promiscuity when using substrates with alternative sugars in the payload: while bare ADPR is a very good substrate, ADP-D-glucose and ADP-L-heptose were not accepted. This discrepancy might stem from constraints in linking the phosphate directly to the sugar ring’s C1’ in these donors, contrasting the out-of-ring C5’-linked ADPR (Table 1, Table S1).

Next, we leveraged another enzymatic route to generate diverse ADDs by using ADP-ribosyl cyclase (ADPRC). As in our previously reported click-ADDing strategy,^30^ which employed this commercial enzyme to generate two ADPR derivatives with bioorthogonal handles, we exploited its controllable base-exchange side reaction with NAD^+^. Using terminal alcohols and N-heterocycles^32–35^ allowed us to synthesize donor candidates with functional groups such as boronic acid or halogens, which we again found to be readily transferred by ApbE (Table 1, Table S1). These results extend the range of functionalities compatible with ApbE-mediated transfer and highlight the advantage of its dependency on ADDs, whose natural abundance means that a variety of biochemical tools can be leveraged to generate diverse donors.

### One-step fluorophore attachment and enzyme immobilization

Having established that ApbE tolerates a diversity of payloads with smaller or similar molecular weight compared to native flavin, we next sought to determine the limits with bulkier molecules. To that end, we could build on our ADPRC-derived click chemistry ADDs (Figure 2A), which constitute a building block allowing to convert virtually any molecule into an ADP-bearing donor candidate via copper(I)-catalyzed azide-alkyne cycloaddition (CuAAC). This strategy reverses our original clickADDing strategy, in which ApbE initially installs a bioorthogonal handle on the target protein and then functionalizes synthetically. Here, we first attached a bulky functional group on the donor and then probed ApbE for direct transfer. We first attached sulfo-Cy3-alkyne to azido-ADPR through CuAAC, after removing excess of azido alcohol from the ADPRC reaction mixture by extraction with ethyl acetate (Figure S1). Gratifyingly, we observed efficient ApbE-mediated conjugation, achieving full conversion within 4 h, as evidenced by a mobility-shifted and intensely fluorescent protein band (Figure 2C).

**Figure 2.**
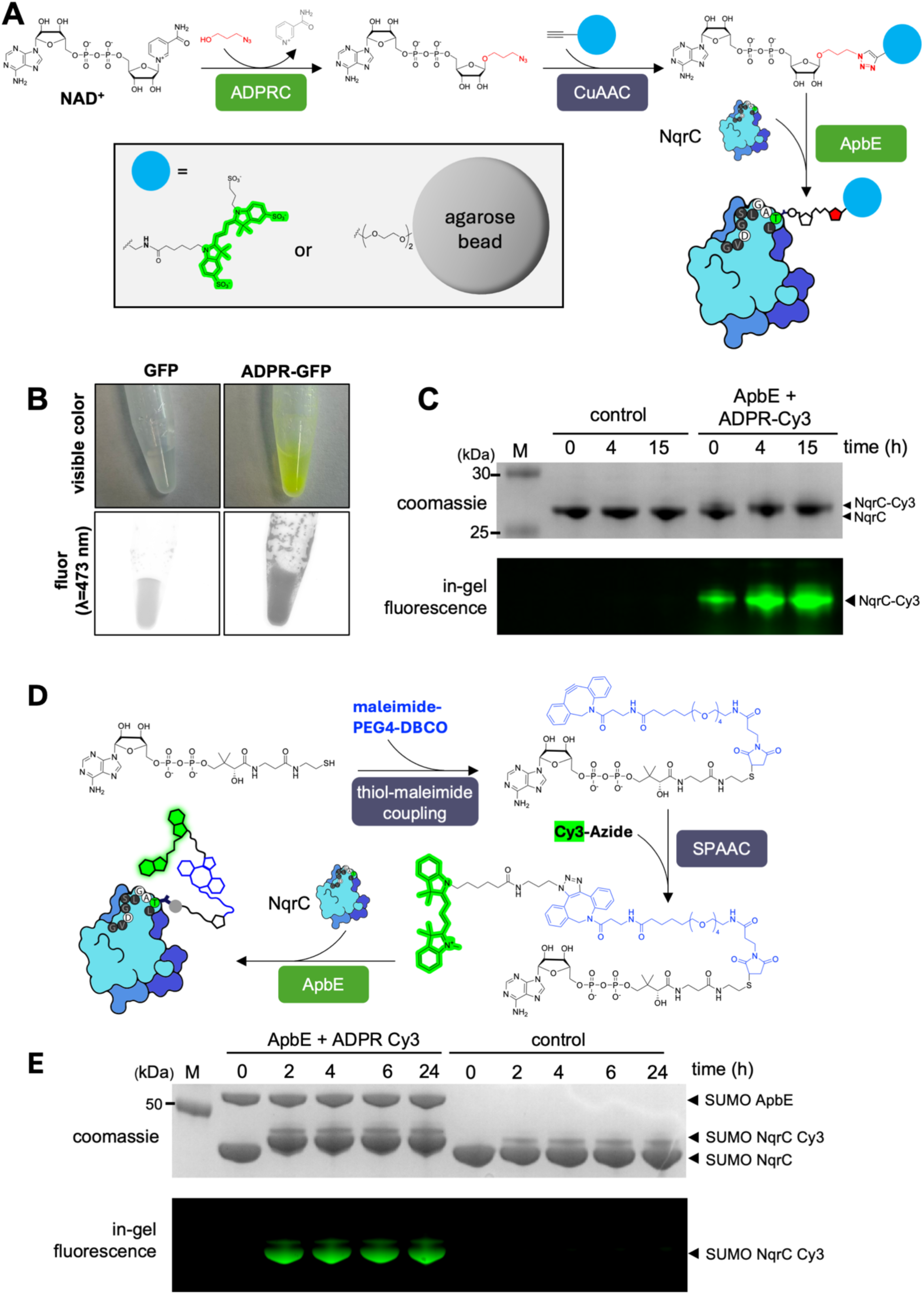
ApbE accommodates bulky cargoes. (A) Overview of the chemoenzymatic strategy used to generate ADPR derivatives by click chemistry. (B) Enzymatic immobilization of GFP by direct, ApbE-mediated protein conjugation to ADPR-functionalized agarose beads. GFP is retained on ADPR-agarose, while an alkynyl-agarose control shows no detectable retention. (C) ApbE-mediated transfer of ADPR-Cy3 enables direct fluorescent labeling of NqrC. SDS-PAGE analysis confirms complete labeling, as indicated by band shift and fluorescence. (D) Workflow of donor generation exploiting the free thiol group of dephospho-CoA (dCoA). dCoA-Cy3 was synthesized via thiol-maleimide coupling and strain-promoted azide-alkyne cycloaddition (SPAAC). (E) ApbE-catalyzed transfer of dCoA-Cy3 and analysis by SDS-PAGE.

At over 1 kDa, substantially larger than FAD (∼786 Da), and negatively charged rather than hydrophobic, this substrate demonstrated that ApbE’s promiscuity extends well beyond modest modifications or truncations of its native donor. Encouraged by this result, we challenged ApbE with an even more extreme substrate: a polymer bead, among the largest payloads employed in bioconjugation. To enable this, we converted commercial alkynyl-agarose to ADPR-agarose by reacting it with purified ADPRC-generated azido-ADPR (Figure 2B). After removing residual copper, we incubated the functionalized resin with an ADD-tag-bearing GFP variant and ApbE for 12 h at room temperature. Astonishingly, we observed efficient conjugation of this payload to GFP, as indicated by the resin’s green color and fluorescence unaffected by extensive washing with elution buffer. In contrast, GFP was almost completely eluted when the same experiment was performed with alkynyl-agarose, confirming that protein retention depended on ADPR-mediated conjugation rather than nonspecific resin binding (Figure 2B). These results show that ApbE accepts donor substrates more than four orders of magnitude bigger than itself and means ADDing can be used for direct, one-step enzymatic protein immobilization.

As an alternative to click chemistry, we also tested whether ApbE would accept bulky ADD donors generated by linking to the free thiol group of dCoA, an approach analogous to the preparation of CoA derivatives for PPTases. We successfully synthesized dephospho-CoA-Cy3 through thiol-maleimide conjugation followed by a strain-promoted azide-alkyne cycloaddition (SPAAC) reaction (Figure 2D). Gratifyingly, subsequent ApbE-catalyzed reactions demonstrated efficient substrate transfer, achieving complete conversion within 4 h, as evidenced by the appearance of a mobility-shifted fluorescent protein band (Figure 2E). These results further demonstrate the versatility of ApbE substrate preparation strategies and the enzyme’s promiscuity toward bulky substrates carrying distinct anchor-linked payloads.

### ApbE accepts FAD-modified peptides and ADP-ribosylated proteins as substrates

So far, we established that ApbE transfers most ADP derivatives regardless of size and demonstrated the convenience of obtaining such donors either commercially directly: as CoA-derivatives that become ADDs with rSAP, as ADPRC-generated ADDs, or as larger ADDs generated via ADPRC-made clickADDs followed by click chemistry, or via maleimide coupling to dCoA. These chemistry first-enzyme second strategies reverse, and thus complement the original ADDing approach (handle conjugation first, chemistry second).

Yet, we also realized that ADDs additionally comprise a large corpus of natural systems, including peptides and proteins. Given ApbE’s activity with large substrates, we therefore next investigated whether this tolerance could be exploited for peptide-protein and protein-protein conjugation. One source of dinucleotide-linked polypeptides are enzymes bearing covalent FAD. Unlike ApbE-mediated FMNylation, these enzymes autocatalytically link various protein side chains to the isoalloxazine ring of the dinucleotide flavin.^36^ As a proof of concept, we thus generated FADylated peptides by tryptic digestion of a representative enzyme, eugenol oxidase (Figure 3A). After heat denaturation, dialysis, and concentration, we incubated the soluble peptide digest with ApbE and the NqrC model acceptor. Analysis by SDS-PAGE revealed indeed the conjugation of different-length peptides to the protein, as indicated by band shift and flavin-caused UV fluorescence. Although the overall reaction efficiency was modest, this unoptimized approach demonstrates the feasibility of using ADDing to attach biopolymers via a natural fluorescent linker to target proteins in a single enzymatic step.

**Figure 3.**
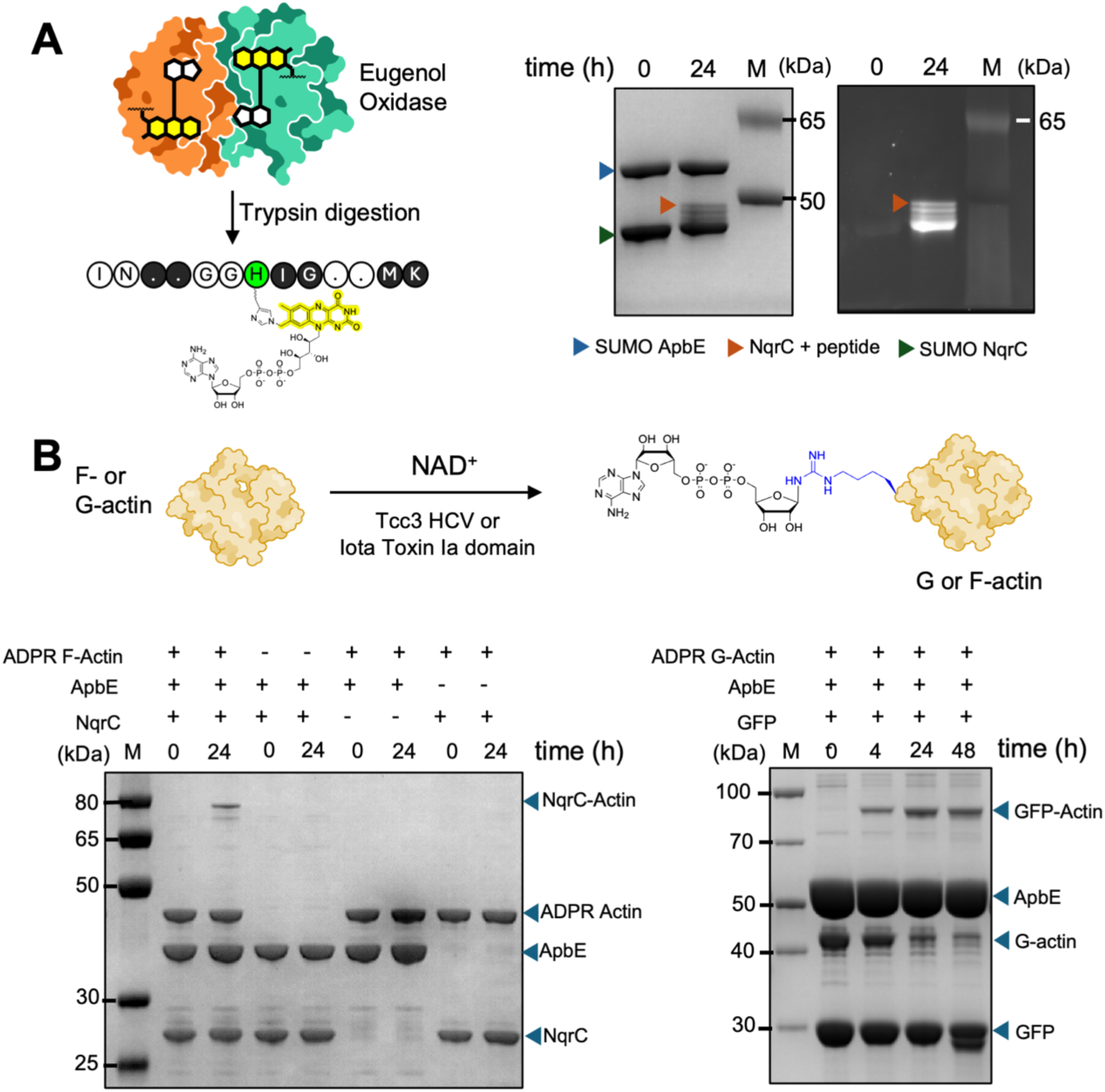
ApbE-mediated conjugation of FADylated peptides and ADP-ribosylated proteins. (A) Overview of FADylated peptide generation (left) and SDS-PAGE analysis (right) showing conjugation of the FADylated peptide to SUMO-NqrC, as indicated by a mobility shift and fluorescence signal. (B) Overview of the enzymatic generation and one-step conjugation of ADP-ribosylated actin to target proteins. SDS-PAGE analysis shows the formation of actin-NqrC conjugates (left) and actin-GFP conjugates (right).

Another, significantly larger source of natural ADD-bearing biopolymers are proteins bearing ADP-ribosylations, a prevalent post-translational modification found in all domains of life.^37–39^ To test whether ADP-ribosylated proteins can act as donors in ApbE reactions, we tested four different systems: ubiquitin, poly(ADP-ribose) polymerase (PARP) 10, and filamentous (F) as well as monomeric globular (G)-actin. We obtained multivalently ADP-ribosylated PARP10 via an autocatalytic self-labelling reaction,^40^ while monovalently modified ubiquitin, F-, and G-actin, were obtained using their dedicated ADP-ribosylase writer enzymes: TccC3 HVR from *Photorhabdus luminescens*,^41^ the Ia domain of *Clostridium perfringens* iota toxin,^42^ and SdeA^519–1100^ from *Legionella pneumophila,*^43^ respectively (Figure S2). Strikingly, we found direct protein-protein conjugation with three out of four of these donors: while ubiquitin did not discernably react, even when we used a small peptide as acceptor (Figure S3), analysis of PARP10 and actin reactions via SDS-PAGE showed a time-dependent formation of a high-molecular weight protein species attributable to the conjugation product. Unsurprisingly, PARP10 reactions did not yield a single defined band, in line with a heterogeneous product, where varying ADP-ribosylation sites lead to a mixture of branched polypeptide products, and where once-transferred PARP10 can again act as donor (Figure S4). In contrast, ApbE-mediated conjugation with actin led to a much more homogenous conjugate and reactions occurred with both molecular forms (Figure 3, S5). Notably, the conversion efficiency for ADP-ribosylated F-actin (approximately 10-20%) and ADP-ribosylated G-actin (∼25%) were comparable to protein-protein conjugations via click-chemistry (Figure 3, S5 and S6). Analysis of the protein-GFP conjugates via semi-native SDS-PAGE (non-reducing and non-denaturing conditions) resulted in a band close to the gel’s pocket for both G-actin-GFP and PARP10-GFP, indicating relatively rapid protein aggregation (Figure S4 and S7). For applications employing this strategy, further optimizations would thus likely require an optimized peptide linker on the acceptor side, serving as a ‘stable landing zone’. Together, these results demonstrate that ApbE can in principle accommodate diverse ADP-bearing polypeptide donor substrates obtained from biochemical sources.

## Discussion

Although numerous biochemical site-specific bioconjugation methods are available, none performs optimal across all relevant criteria. To some extent, this diversity is an inherent strength of the field: hydrophobic or hydrophilic linkers, reversible or irreversible attachments, strict motif specificity or relaxed acceptor profiles represent alternative properties whose desirability depends on the application. Other criteria, including ease of enzyme production, donor accessibility, and donor promiscuity, more clearly distinguish strengths from limitations. Here we have shown that bioconjugation with ApbE is particularly attractive where the latter properties matter. As evident from the multiple donor-generation strategies described in this work, ADP-moieties are abundant in nature, and an extensive biochemical toolbox for installing and modifying these scaffolds complements the commercial availability of diverse ADD building blocks. Notably, ADDing also benefits from the maturity of bioconjugations using PPTases like Sfp, since dephosphorylation of CoA derivatives yields ADD-tag-compatible substrates. Indeed, ADDing can be considered a generalization of PPTase-based labelling regarding donors: despite being structurally unrelated, both enzymes install payloads as O-linked phosphodiesters, but ADDing acts on donors without the pantetheine linker as much as those that contain it. Beyond this broader donor scope, ADDing shares a key strength of PPTases: remarkable tolerance to payload size, demonstrated here by the efficient conjugation of ADP-functionalized polymer beads and ADP-ribosylated proteins. Whereas this size tolerance determines the range of accessible conjugation applications, donor availability primarily influences their practicality and cost. Together, these properties make direct conjugation by ADDing an attractive alternative to our previously described two-step strategy, in which an enzymatically installed handle is subsequently functionalized by click chemistry.^30^ This reversal of the workflow—chemistry first, enzymatic conjugation second—should be particularly advantageous when chemical modification of proteins or complex biological samples presents a challenge, and when bioorthogonal chemistry is not an option.

Although our results reveal ApbE’s exceptionally broad substrate tolerance, its promiscuity is not unrestricted. Besides a stringent requirement for the ADP anchor, our donor profiling shows that substrates bearing certain substituents directly linked to the payload phosphate, such as glucose, were not accepted. Since ribose at the same position (albeit linked via an out-of-ring carbon) was efficiently transferred, it appears that the size, geometry or flexibility of the substituent adjacent to the phosphate can influence productive substrate binding or catalysis.

Another curiosity is ApbE’s inactivity toward NAD⁺, while being active on NADH. As these molecules only differ in their hydrogenation and thus charge at the nicotinamide ring, these properties appear to influence acceptance. The mechanistic basis and whether this might affect a broader donor molecule class remains to be investigated. Despite these exceptions, our work shows that, generally, the majority of payloads is compatible with direct ADDing. The range of functionalities we demonstrated illustrates the potential for both fundamental research and biotechnology. ADDing can functionalize proteins with small molecules to probe molecular interactions, visualize biological processes, or introduce affinity handles, such as nitro groups, boronic acids, free thiols, or halogens, as shown in this work, in addition to the azide and alkyne handles from our previous approach. Large payloads such as peptides and proteins may be applied for biomolecular fusions inaccessible by genetic manipulation and might prove particularly useful for biomedical applications including multispecific antibody formats.

A particularly attractive extension of ApbE-mediated conjugation is its compatibility with ADP-ribosylated proteins, opening new possibilities for harnessing this biologically important post-translational modification for protein bioconjugation. ADP-ribosylation is a widespread and important PTM involved in diverse cellular processes, including DNA damage responses, cellular signalling and host-pathogen interactions,^38,39,44^ research areas that might also directly benefit from the ability of ApbE to attach such biomolecules to suitable acceptors. For example, this approach could provide a means of conjugating functionalities to natural or deliberately ADP-ribosylated proteins, and enable their enrichment or immobilization for biochemical studies.

Several aspects make ADDing an attractive alternative to established methods for site-specific protein-protein conjugation. Widely used approaches such as Sortase A-mediated ligation and protein trans-splicing predominantly target protein termini,^6^ whereas SpyTag/SpyCatcher allows more flexible attachment sites but introduces a comparatively large protein domain into the conjugate.^45^ ADDing instead combines a short acceptor motif compatible with internal protein loops, with the ability to directly attach bulky cargo without additional protein domains. In this respect, it resembles the more recently developed lysine acylation using conjugating enzymes (LACE), which enables site-specific attachment of proteins through a minimal four-residue recognition motif. However, LACE requires chemically activated protein thioesters, and its demonstrated protein donor scope remains largely restricted to ubiquitin and related proteins.^46^ In contrast, ADDing can exploit enzymatic ADP-ribosylation to generate protein donors, potentially accommodating a much broader range of cargo proteins. The PPTase Sfp likewise combines short acceptor tags with substantial donor promiscuity, but requires CoA-derived substrates carrying a phosphopantetheine linker.^8,22^ ADDing thus offers a complementary combination of flexible acceptor placement, broad payload tolerance, and access to diverse donor substrates through biochemical or chemoenzymatic routes.

Compared with handle-based two-step approaches, including our own,^25,47–49^ direct conjugation of bulky payloads can circumvent several limitations, including additional non-native linkers and procedural complexity. Perhaps more importantly, it reverses the order of chemical and enzymatic modification. In conventional enzyme-first strategies, the chemical reaction occurs on the target protein, imposing constraints on both the biological sample and the applicable chemistry. In contrast, chemistry-first, enzyme-second approaches allow the payload to be prepared independently, potentially exploiting a much broader repertoire of synthetic transformations before enzymatic conjugation under mild, biologically compatible conditions. In the case of ADDing, this produces a phosphodiester linkage resembling naturally occurring PTMs, which may offer advantages in biological compatibility and warrants investigation of its immunogenicity. Moreover, the highly hydrophilic nature of this linkage could prove particularly attractive for antibody-drug conjugates, where linker hydrophilicity can help counteract payload-induced aggregation and unfavorable pharmacokinetics.

In sum, we here describe ADDing via ApbE as a powerful bioconjugation platform offering multiple features that make it attractive for site-specific protein modification. Its recognition of a short, seven-residue peptide motif with several positions tolerant to exchanges facilitates flexible incorporation into target proteins.^50,51^ We have shown that the enzyme accommodates substantial variation in the transferred moiety, including bulky payloads, and its substrates can be accessed through an arsenal of chemical and chemoenzymatic synthesis routes.^52^ Although this study explores only a subset of possible substrate-preparation strategies, the chemistry of a set of natural dinucleotides including NAD, dephospho-CoA, and ADPR as well as the synthetic chemically addressable derivatives offer considerable scope for diversification. The combination of ApbE’s substrate promiscuity, the diversity of naturally occurring substrates, and established chemoenzymatic synthetic routes enables the transferred functionality to be tailored to specific applications without requiring a new enzyme variant for each cargo.

## Ethics declarations

