## Supplementary Information for "Direct Bioconjugations of Diverse Payloads using an Extraordinarily Donor-Promiscuous Flavin Transferase"

### 1. Supplementary Figures

#### 1.1. Purification of Azido ADPR

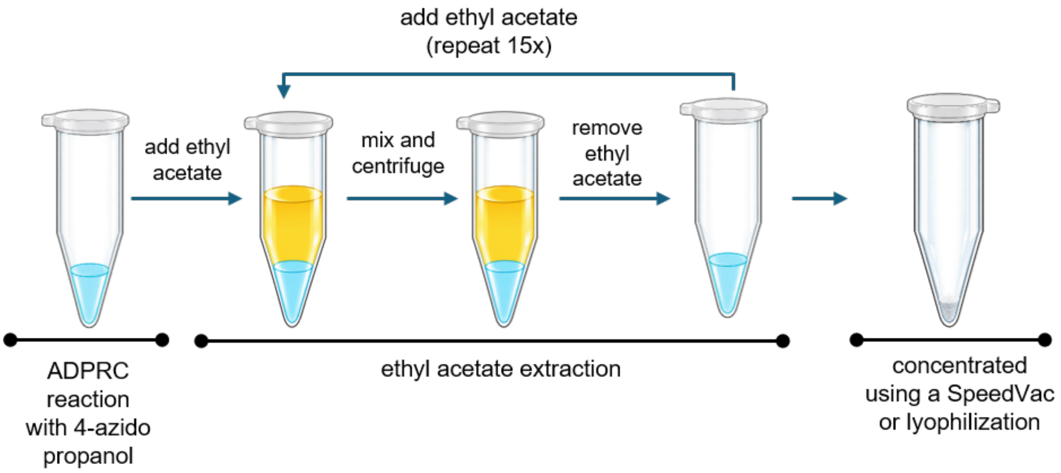


**Figure S1.** Ethyl acetate extraction protocol for removing excess 3-azidopropanol following ADPRC transglycosylation reactions.

#### 1.2. Deconvoluted Mass Spectra of ADP-ribosylated G-actin and F-actin

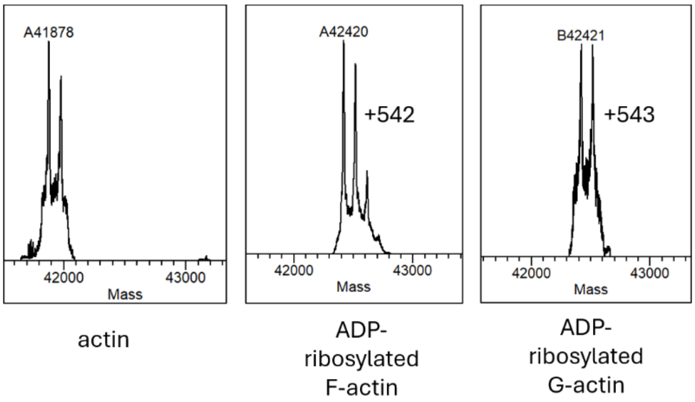


**Figure S2.** Deconvoluted Mass Spectra of ADP-ribosylated G-actin and F-actin.

#### 1.3. ApbE reaction of ADPR-ubiquitin with F2-peptide

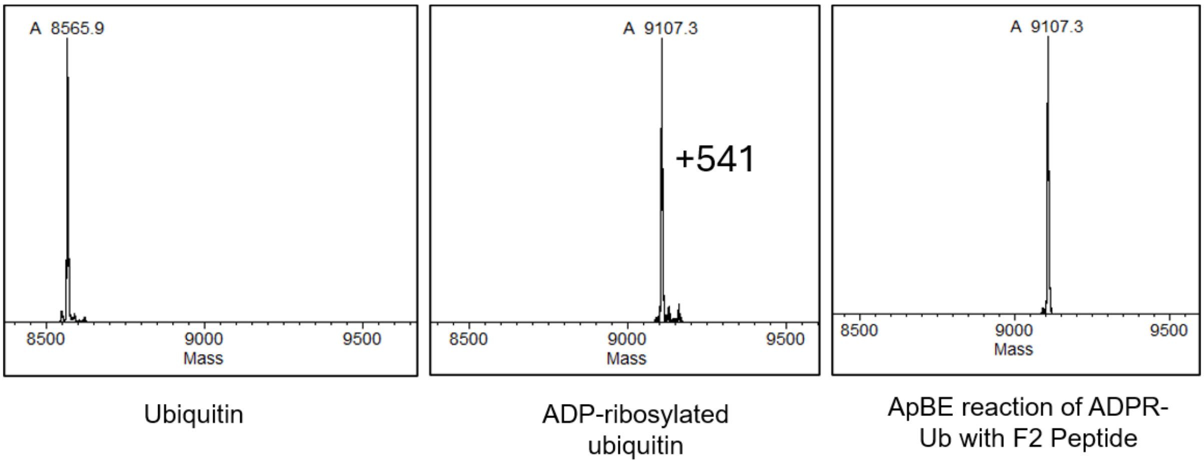


**Figure S3.** Deconvoluted mass spectrum of ubiquitin ADP-ribosylation by SdeA$^{519\text{-}1100}$ and unsuccessful F1 peptide (GVDGLSGATLTSGG) conjugation to ADPR-ubiquitin.

#### 1.4. ApbE reactions of multi-ADPR-PARP10 with NqrC and GFP

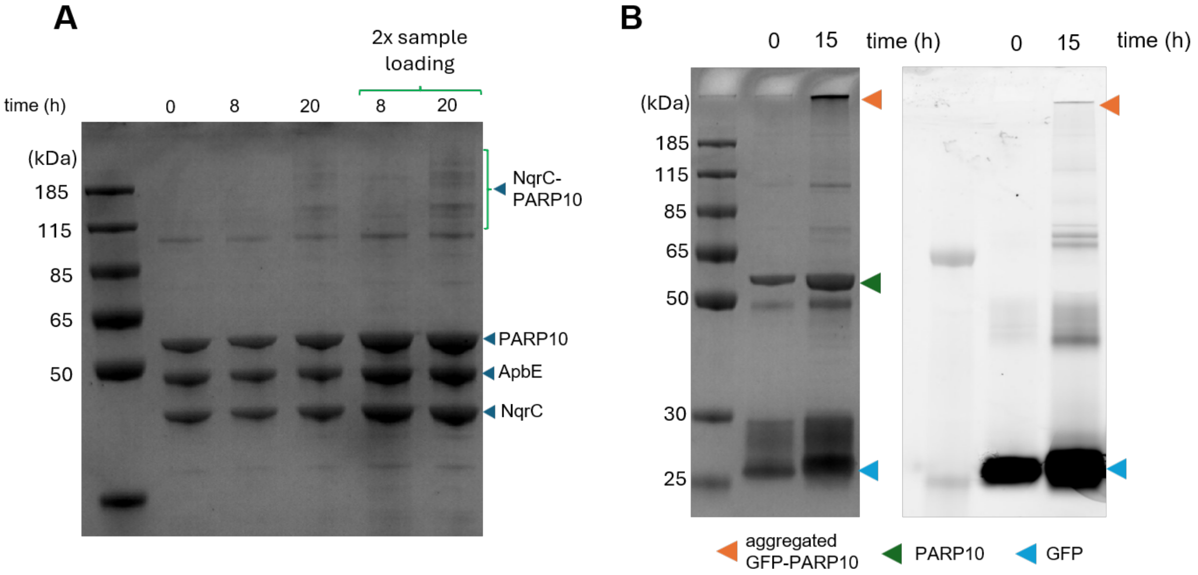


**Figure S4.** (A) SDS–PAGE analysis of the ApbE-catalyzed reaction between multi-ADP-ribosylated PARP10 and NqrC. The appearance of a new band above 115 kDa shows the attachment of NqrC to multi-ADP-ribosylated PARP10. (B) Non-denaturing SDS–PAGE analysis of the ApbE-catalyzed reaction between ADP-ribosylated PARP10 and GFP. The retention of the resulting PARP10–GFP conjugate at the very top of the gel suggests protein aggregation.

##### 1.5. Optimization of F-Actin NqrC Conjugation

| Conditions | protein concentrations (mg/ml) | | | |
|---|---|---|---|---|
| | [SUMO ApbE] | [ApbE] | [NqrC] | [ADPR-F1 actin] |
| **A** | - | - | 3 | 5 |
| **B** | 10 | - | 3 | 5 |
| **C** | - | 10 | 3 | 5 |
| **D** | 10 | | 3 | 5 |
| **E** | 5 | - | 3 | 5 |
| **F** | 5 | - | - | 5 |

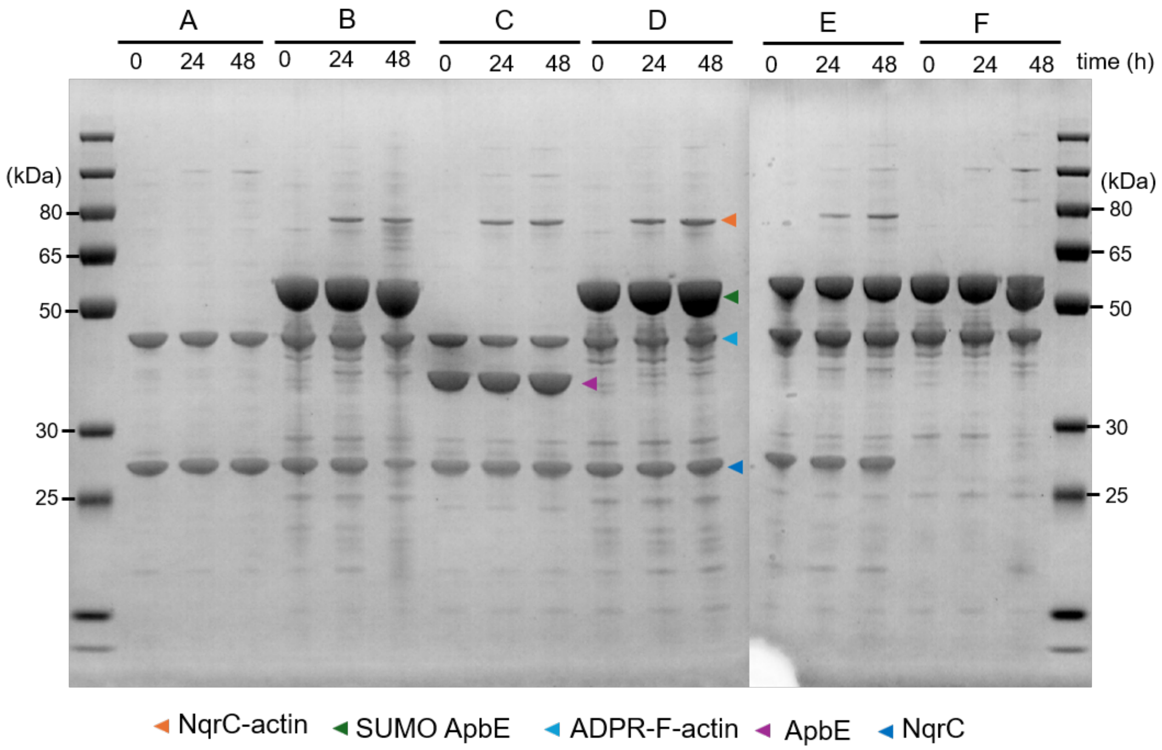


**Figure S5.** Optimization of reactions conditions for ApbE reaction with ADP-ribosylated F-actin and NqrC.

#### 1.6. ApbE reaction of monomeric ADPR-G-actin with NqrC

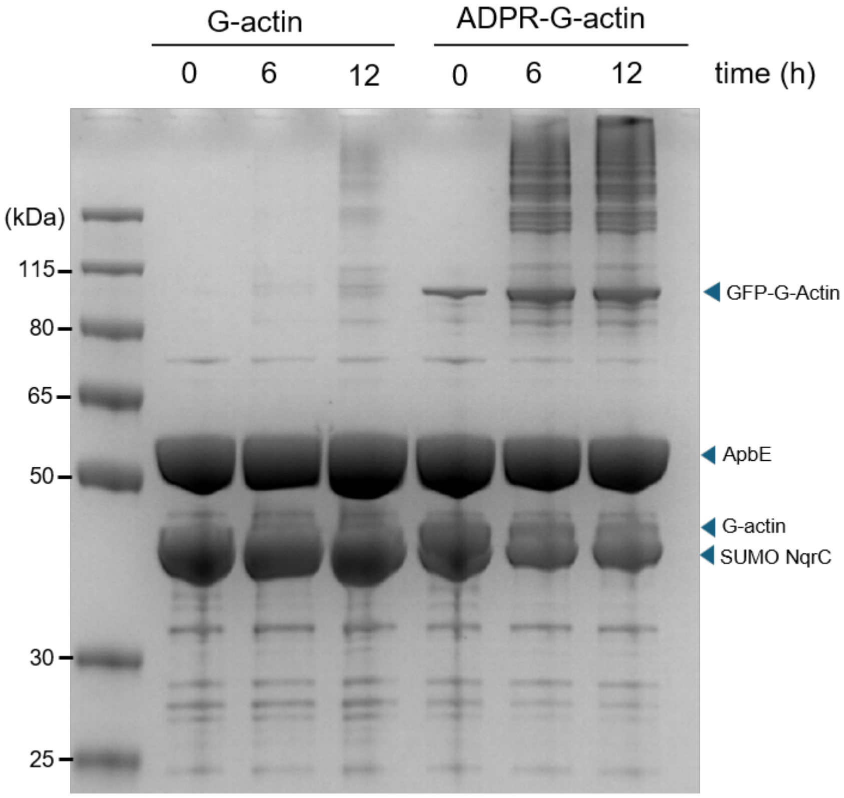


**Figure S6.** SDS–PAGE analysis of the ApbE-catalyzed reaction between monomeric ADP-ribosylated G-actin and NqrC. The appearance of a new band at approximately 90 kDa suggests the formation of G-actin–NqrC conjugates.

#### 1.7. Non-denaturing gel of ApbE reaction of G-actin with GF

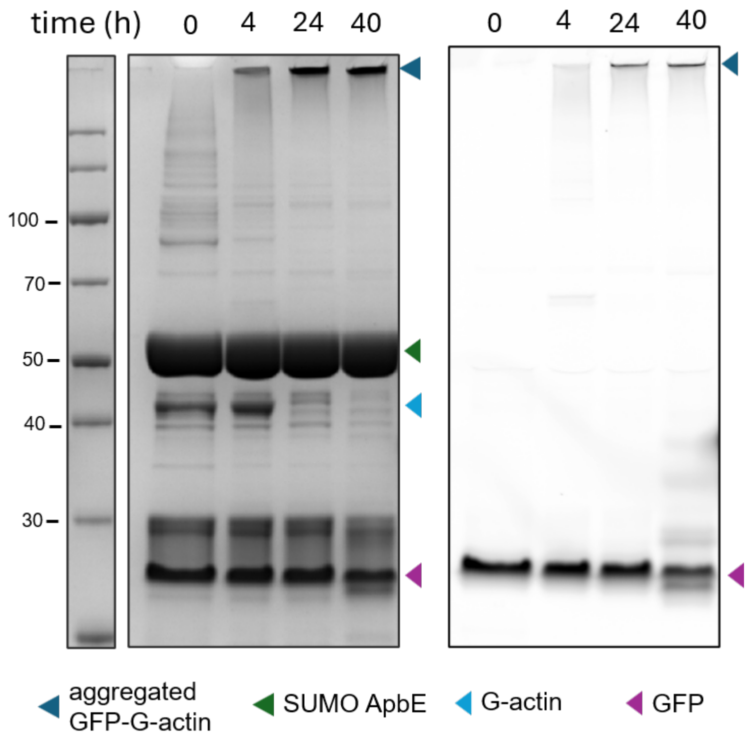


**Figure S7.** Non-denaturing SDS–PAGE analysis of the ApbE-catalyzed reaction between mono-ADP-ribosylated G-actin and GFP. The samples are the same as those analysed in Figure 3C. The accumulation of the resulting G-actin–GFP conjugate at the top of the gel suggests protein aggregation.

### 2. Supplementary Tables

#### 2.1. Deconvoluted Mass Spectra of ADDs Screening Results

**Table S1.** Complete deconvoluted mass spectra of ADDs screening results.

<table>
<tr><th colspan="2">A. Control SUMO NqrC</th><th colspan="2">B. SUMO NqrC + AppG</th></tr>
<tr><td rowspan="3">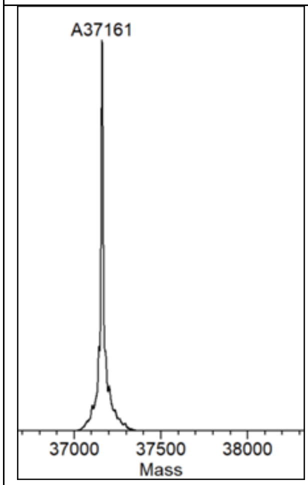
</td><td>Exp. Mass (Da): 37161</td><td rowspan="3">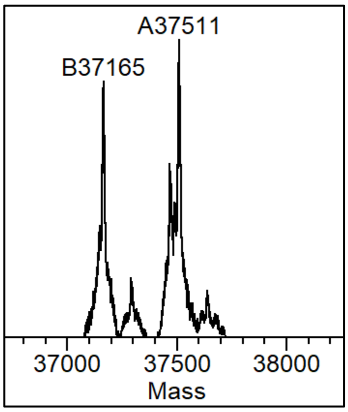
</td><td>Exp. Mass (Da): 37507</td></tr>
<tr><td>Obs. Mass (Da): 37161</td><td>Obs. Mas (Da): 37511</td></tr>
<tr><td>Notes: -</td><td>Notes:<br><br>Exp. Mass = S.NqrC (37161) + AMP (363.2) – OH (17) = 37507 (Da)</td></tr>
<tr><th colspan="2">C. SUMO NqrC + ADPR</th><th colspan="2">D. SUMO NqrC + ADPR pNP</th></tr>
<tr><td rowspan="3">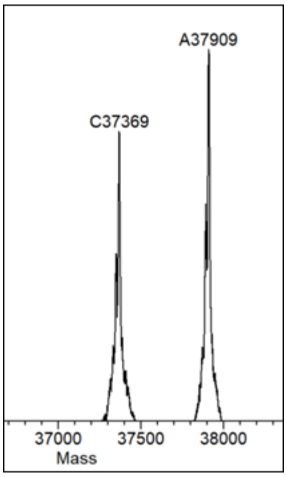
</td><td>Exp. Mass (Da): 37374</td><td rowspan="3">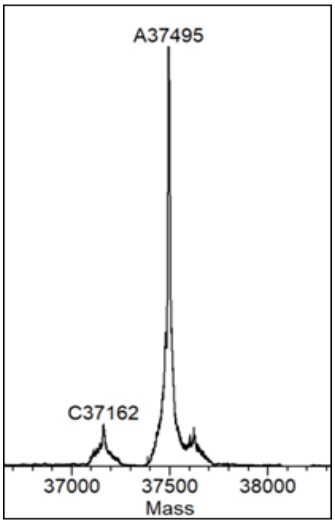
</td><td>Exp. Mass (Da): 37477</td></tr>
<tr><td>Obs Mass (Da): 37369, 37909</td><td>Obs Mass (Da): 37495</td></tr>
<tr><td>Notes:<br><br>Exp. Mass = S.NqrC (37161) + phosphoribose (230) – OH (17) = 37374 (Da). Additional mass of 37909 (37374 + 540) because of additional mass of ADPR caused by SUMO non enzymatic ADP-ribosylation.</td><td>Notes:<br><br>Exp. Mass = 37161 (S.NqrC) + (333.2) – OH (17) = 37477 (Da). The additional 17 Da might be due to hydroxylation</td></tr>
</table>

##### E. SUMO NqrC + NADH

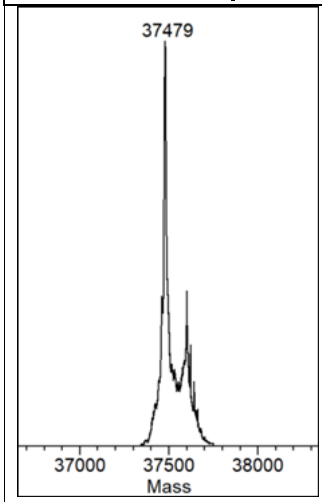


Exp. Mass (Da): 37478

Obs. Mass (Da): 37479

**Notes:**

Exp. Mass = 37161 (S.NqrC) + NMN (334) – OH (17) = 37478(Da)

##### F. SUMO NqrC + NAD$^+$

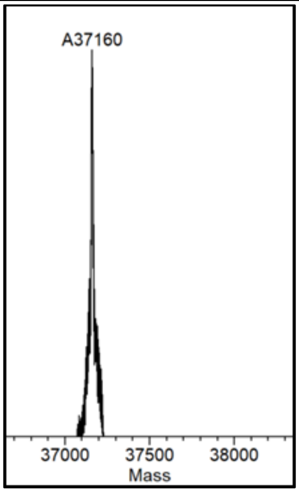


Exp. Mass (Da): 37478

Obs. Mass (Da): 37161

**Notes:**

Exp. Mass = 37161 (S.NqrC) + NMN (334) – OH (17) = 37478(Da)

##### G. SUMO NqrC + ADP-Glucose

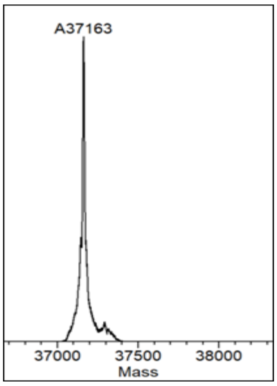


Exp. Mass (Da): 37403

Obs. Mass (Da): 37161

**Notes:** Exp. Mass = 37161 (S.NqrC) + phospho glucose (259) – OH (17) = 37403 (Da)

##### H. SUMO NqrC + ADP-L-Heptose

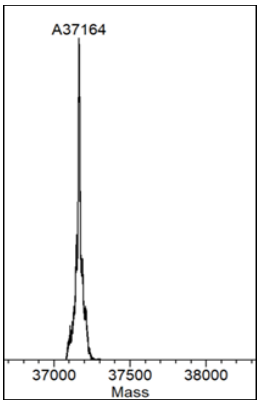


Exp. Mass (Da): 37433

Obs. Mass (Da): 37161

**Notes:** Exp. Mass = 37161 (S.NqrC) + phosphor-L-heptose (289) – OH (17) = 37433 (Da)

##### I. SUMO NqrC + Dephospho-Acetyl-CoA

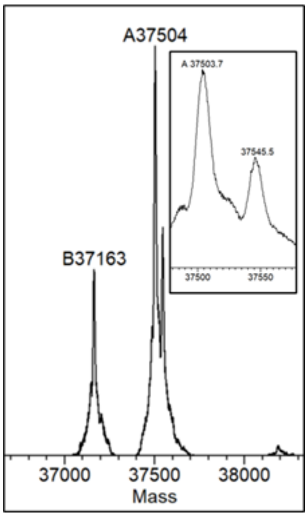


Exp. Mass (Da): 37544

Obs. Mass (Da): 37545.5, 37504

Notes: Exp. mass = (S.NqrC) 37161 + acetyl PPT (400) – OH (17) = 37544(Da). The loss of acetyl mass is most likely due to thioester hydrolysis by acidic buffer or in source fragmentation.

##### J. SUMO NqrC + Dephospho-Lauryl-CoA

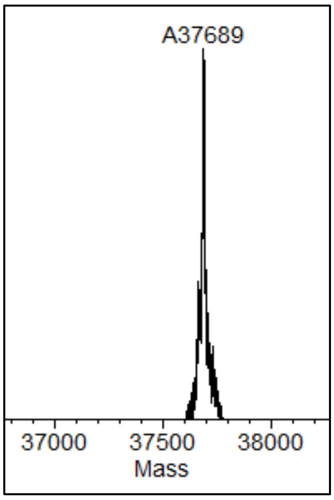


Exp. Mass (Da): 37694

Obs. Mass (Da): 37157

Notes: Exp. Mass = 37161 (S.NqrC) + lauryl PPT (540) – OH (17) = 37694(Da)

**K. SUMO NqrC + Dephospho-CoA**

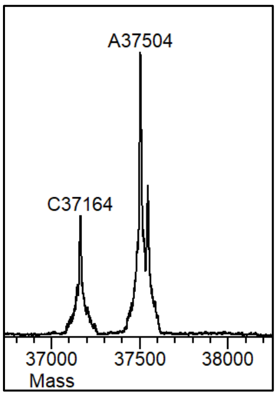


Exp. Mass (Da): 37502.3

Obs. Mass (Da): 37504

**Notes:** Exp. mass = (S.NqrC) 37161 + PPT (358.3) – OH (17) = 37502.3 (Da).

**L. SUMO NqrC + 6-bromo-isoquinoline ADPR**

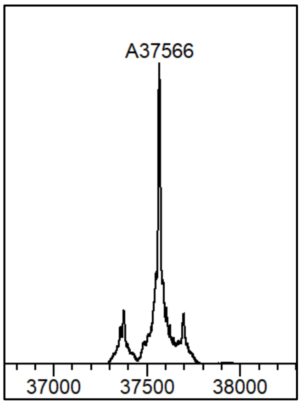


Exp. Mass (Da): 37566

Obs. Mass (Da): 37566

**Notes:** Exp. Mass = 37161 (S.NqrC) + phosphoribose 6-bromo-isoquinoline (422) – OH (17) = 37566 (Da)

**M. Control F1-MBP**

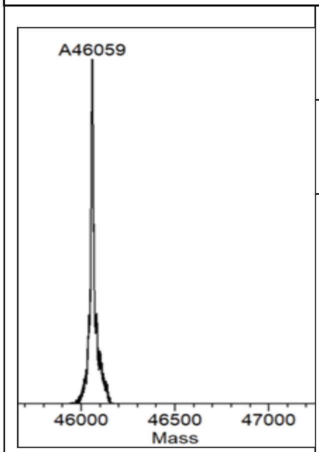


Exp. Mass (Da): 46059

Obs. Mass (Da): 46059

**Notes: -**

**N. F1-MBP + NADPH**

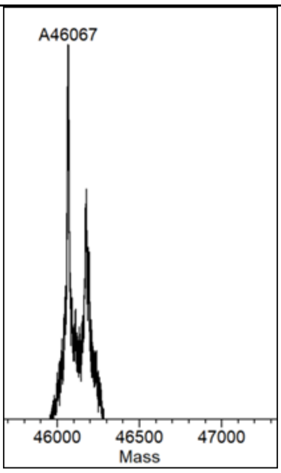


Exp. Mass (Da): 37161

Obs. Mass (Da): 37161

**Notes:**

Exp. Mass = 46065 (F1-MBP) + + NMN (334) – OH (17) = 37478(Da)

= 46382(Da).

**O. F1-MBP + Acetyl-CoA**

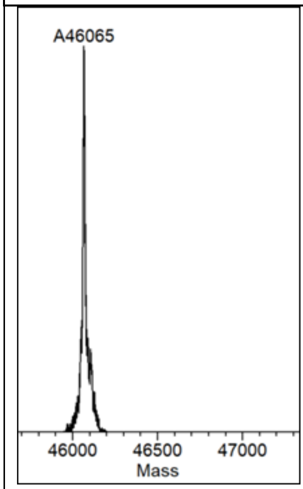


Exp. Mass (Da): 46376

Obs. Mass (Da): 46065

**Notes:**

Exp. Mass = 46065 (F1-MBP) + NMN (334) – OH (17) = 46376 (Da).

**P. F1-MBP + 3-phenylboronic ADPR**

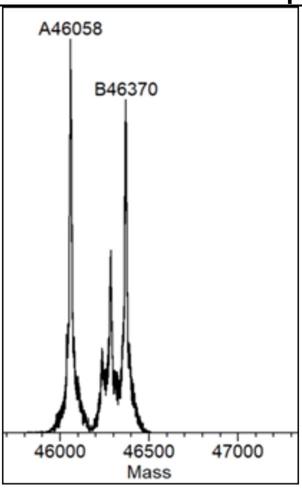


Exp. Mass (Da): 46058

Obs. Mass (Da): 46404

**Notes:** Exp. Mass = 46058 (F1-MBP) + phosphor-ribosyl phenyl-boronic acid (363) – OH (17) = 46404(Da). The loss of 34 Da corresponds to the fragmentation of 2 hydroxyl groups.

| **Q. MBP-F1 + Dephospho palmitoyl CoA** | | **R. F1-MBP + Imidazole** | |
|---|---|---|---|
| 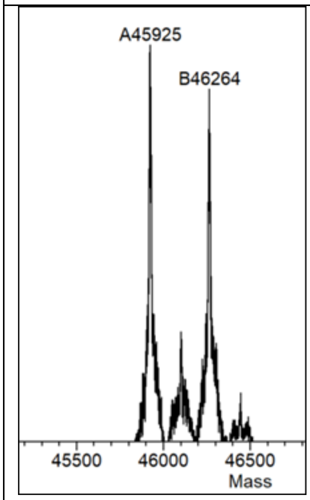 | Exp. Mass (Da): 46466 | 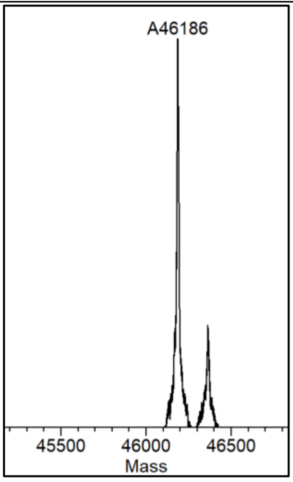 | Exp. Mass (Da): 46186 |
| | Obs. Mass (Da): 46264 | | Obs. Mass (Da): 46186 |
| | **Notes:** Exp. Mass = 45922 (MBP-F1) + palmitoyl-PPT (561) – OH (17) = 37705 (Da). The mass of 46264 Da corresponds to the mass of S.NqrC + PTT. The loss of palmitoyl moiety is most likely due to thioester hydrolysis by acidic buffer or in source fragmentation. | | **Notes:** Exp. Mass = 45922 (MBP-F1) + phospho-ribose ADPR (281) – OH (17) = 46186 (Da). |

#### 3. Materials and Methods

##### 3.1. General considerations

All standard chemicals were obtained from commercial suppliers including Sigma-Aldrich, TCI, and Thermo Fisher Scientific. Panel of ADDs substrates was purchased from Jena Biosciences (Gp2G, Ap3G, ADPR pNP), MedChem (ADPR), Merck (Acetyl CoA, Lauroyl COA), Roth (NAD, NADH), Oriental Yeast (NADPH), InvivoGen (ADP-L-Heptose), and TargetMol (dephospho CoA). Other specialized reagent such as 3-azidopropanol and alkynyl agarose resin were purchased from Sigma-Aldrich. Alkynyl-sulfo-Cy3 and Azido-Cy3 was sourced from Jena Biosciences. ADP-ribosyl cyclase (ADPRC) from *Aplysia californica* was acquired from Sigma-Aldrich. rhUbiquitin was purchased from Bio-techne. The EZ-Link™ Maleimide-PEG4-DBCO was purchased from ThermoFisher Scientific.

PCR and molecular biology applications were purchased from New England Biolabs (NEB). Oligonucleotides and primers were obtained from Integrated DNA Technologies (IDT), and plasmid and DNA purification was performed using QIAGEN kits. Protein purification was carried out using Ni Sepharose 6 Fast Flow resin (Cytiva), and buffer exchange was performed using PD-10 desalting columns (Cytiva). Protein identity and purity were confirmed by electrospray ionization mass spectrometry (ESI-MS) using a Waters Acquity H-Class UPLC system coupled to a Waters Xevo G2 QTOF mass spectrometer. Coomassie-stained SDS–PAGE gels were imaged using a Bio-Rad GelDoc system, and fluorescence imaging was conducted on a GE Typhoon FLA 9500 scanner.

##### 3.2. Plasmid construction

All plasmids used in this paper was obtained from previous studies.[1–6]

##### 3.3. Protein expression and purification

**Expression and purification of apo ApbE**

Apo ApbE was expressed as a SUMO fusion from the pBAD SUMO ApbE plasmid in a riboflavin-deficient *E. coli* strain (BL21 AI *ΔRibB*). An overnight culture was diluted into LB medium supplemented with 50 µg/mL ampicillin and 20 µg/mL riboflavin and grown to an $OD_{600}$ of ~0.8. Cells were washed three times with LB medium to remove residual riboflavin, resuspended in riboflavin-free LB. After the final wash, the cells were kept on ice for one hour as a starvation step to further reduce the intracellular flavin content, and then induced with 0.02% (w/v) arabinose. Cultures were incubated at 17 °C with shaking (200 rpm) for 72 h before harvesting by centrifugation at 6,000 rpm for 30 min. Protein purification followed previously reported protocols. Cell pellets were resuspended in 50 mM Tris-HCl (pH 7.5) containing 0.5 mM PMSF and lysed by sonication (5 s on, 10 s off, 10 min total). Clarified lysates were obtained by centrifugation at 11,000 rpm for 60 min and filtered through a 20 µm filter prior to loading onto a gravity-flow Ni-NTA column. After binding at 4 °C for 30 min, the column was washed with 50 mM Tris-HCl (pH 7.5) containing 20 mM imidazole, and ApbE was eluted with 300 mM imidazole. The eluted protein was desalted into 50 mM Tris-HCl (pH 7.5), aliquoted, flash-frozen, and stored at −80 °C.

**Expression and purification of NqrC**

SUMO-fused NqrC was expressed from the pBAD SUMO NqrC construct in *E. coli* NEB10β cells. Cultures were induced at an $OD_{600}$ of ~0.8 with 1 mM IPTG and incubated at 24 °C for 24 h. Cells were collected by centrifugation (6,000 rpm, 20 min), and purification was performed using the same Ni-NTA-based protocol described for ApbE.

**Expression and purification of F1-MBP and MBP-F1**

F1-MBP and MBP-F1 constructs, containing N- or C-terminal $His_6$ tags and a 12-amino-acid F1 tag (GVDGLSGATLTS), were expressed from pET28 vectors in *E. coli* BL21(DE3) cells. Protein expression was induced at an $OD_{600}$ of ~0.8 with 1 mM IPTG, followed by incubation at 24 °C for 24 h. Cells were harvested by centrifugation (6,000 rpm, 20 min), and proteins were purified using the same procedure as described for ApbE.

**Expression and purification of GFP–F2**

GFP–F2 (eGFP with F2 tag, GVDGLSGATL) was expressed from a pET28 vector in *E. coli* BL21(DE3) cells grown in TB medium containing 50 µg/mL ampicillin at 24 °C with shaking (200 rpm) for 16 h. Cells were harvested by centrifugation (4,500 rpm, 30 min) and resuspended in 20 mM Tris-HCl (pH 8.0) supplemented with 500 mM NaCl and 0.5 mM PMSF. Following sonication, lysates were subjected to heat treatment at 60 °C for 30 min and clarified by centrifugation at 11,000 rpm for 60 min. The supernatant was filtered and applied to a Ni-NTA gravity column. After washing with 20 mM imidazole, eGFP–F2 was eluted with 500 mM imidazole, desalted into 20 mM Tris-HCl (pH 8.0), 150 mM NaCl, aliquoted, flash-frozen, and stored at −80 °C.

**Expression and purification of Iota Toxin Ia Domain**

Iota toxin Ia domain fused with MBP was expressed using a pET28 vector in *E. coli* BL21(DE3) cells grown in TB medium containing 50 µg/mL ampicillin until OD600 reached 0.6. After induction with 1 mM IPTG, cells were grown at 24 °C overnight with shaking (200 rpm) for 16 h. For purification, cells were harvested by centrifugation and resuspended in lysis buffer (20 mM sodium phosphate, pH 7.4, 500 mM NaCl and 5 mM imidazole) supplemented with 1 mM phenylmethylsulfonyl fluoride (PMSF) and a protease inhibitor cocktail. Cells were lysed by sonication, and the lysate was clarified by centrifugation. The resulting supernatant was loaded onto a Ni-NTA resin column. The column was washed sequentially with lysis buffer and washing buffer (lysis buffer supplemented with 30 mM imidazole). Ia was eluted with the same buffer supplemented with 500 mM imidazole and subsequently desalted into 20 mM Tris-HCl (pH 7.4) containing 50 mM NaCl. The purified protein was flash-frozen, and stored at −80 °C.

**Expression and purification of TccC3 HVR**

The ADP-ribosyltransferase domain of Photorhabdus luminescens TccC3 (TcART) was expressed as an MBP fusion in *E. coli* BL21-CodonPlus(DE3)-RIPL cells carrying plasmid pB656, as described previously.[2] Protein expression was induced with 0.1 mM IPTG overnight at 22°C. Cells were lysed in 20 mM Tris-HCl pH 8.0, 150 mM NaCl, and the recombinant protein was purified by Ni-IDA affinity chromatography. After elution with 250 mM imidazole, the protein was dialyzed against 20 mM Tris-HCl pH 8.0, 150 mM NaCl and stored at -20°C.

**Expression and purification of PARP10**

The GST-tagged catalytic domain of PARP10 (amino acids 818-1025) was purified as described before.[3]

**Expression and Purification of SdeA**

SdeA was expressed as an N-terminal $His_6$-SUMO fusion from a pET28-based expression plasmid. For protein expression, an overnight culture of *E. coli* BL21(DE3) was inoculated into 400 ml of LB medium supplemented with ampicillin. Cultures were grown to an OD600 of 0.6–0.8, induced with 0.2 mM isopropyl β-D-1-thiogalactopyranoside (IPTG) and incubated at 18 °C with shaking for 16–18 h. Cells were harvested by centrifugation at 6000 × g, resuspended in 20 mM Tris-HCl, pH 8.0, 100 mM NaCl, and lysed by sonication in the presence of protease inhibitors cocktails. The soluble fraction was collected by centrifugation at 12,000 × g at 4 °C for 60 minutes. $His_6$-SUMO-$SdeA^{519-1100}$ was purified using NiNTA beads (Qiagen). The bound protein was eluted in lysis buffer + 300 mM imidazole. The purified protein was desalted into 25 mM Tris-HCl (pH 7.5) containing 150 mM NaCl, 5% glycerol and 1 mM dithiothreitol (DTT), flash-frozen, and stored at −80 °C.

**Purification of rabbit skeletal muscle actin**

Rabbit skeletal muscle α-actin was purified from muscle acetone powder (Pel-Freez Biologicals) as previously described.[1,7] Briefly, actin was extracted in G-buffer (5 mM Tris-HCl pH 7.5, 0.2 mM $CaCl_2$, 0.5 mM ATP, 1 mM DTT) and purified through a polymerization-depolymerization cycle . Polymerization was induced by the addition of 2 mM $MgCl_2$ and 100 mM KCl. After high-salt treatment (800 mM KCl) to remove actin-binding proteins, filaments were collected by ultracentrifugation and depolymerized by dialysis against G-buffer. Purified G-actin was aliquoted, flash-frozen in liquid nitrogen and stored at -70°C.

**3.4. Screening of ADDs derivatives**

The ApbE reaction for activity screening was performed in 20 ul reaction volume containing 1 mg/ml ApbE, 3 mg/ml target proteins, 1 mM substrates (ADD derivatives), 1 mM EDTA, and 10 mM $MgCl_2$ in 50 mM Tris-HCl (pH 8.0). Reactions were incubated at 30 °C for 3–12 h.

For semiquantitiave analysis, the successful conjugation was detected using ESI-MS analysis. using a Waters Acquity H-Class UPLC system coupled to a Waters Xevo G2 QTOF mass spectrometer equipped with a photodiode array (PDA) detector. Protein separations were performed on an Acquity BEH C4 column (150 × 2.1 mm, 1.7 µm; Waters) maintained at 40 °C. The mobile phase consisted of 0.1% (v/v) formic acid in water (solvent A) and acetonitrile (solvent B), delivered at a flow rate of 0.3 mL/min. The chromatographic gradient began at 5% solvent B, increased linearly to 70% over 10 min, followed by a ramp to 95% solvent B between 10 and 13 min, and was then returned to 5% solvent B by 20 min to allow column re-equilibration. Protein samples were diluted to final concentrations of 3–5 µM and analyzed in positive-ion ESI mode with an injection volume of 5 μL. Data were acquired and processed using MassLynx software. Charge-state distributions were deconvoluted using MagTran.

##### 3.5. ADPRC reaction to make 4-phenyl boronic acid ADPR

4-Phenylboronic acid ADP-ribose was synthesized using *A. californica* ADP-ribosyl cyclase (ADPRC; Sigma-Aldrich) with $NAD^+$ as the substrate. Reactions were carried out in 50 mM HEPES buffer (pH 7.0) supplemented with 5 mM $MgCl_2$. Each reaction contained 10 mM $NAD^+$, 2 μM ADPRC, and 2 mM 3-(Hydroxymethyl)phenylboronic acid. Reaction mixtures were incubated at 37 °C for 1 h and terminated by heat inactivation of ADPRC at 95 °C for 10 min.

##### 3.6. One-pot ADPRC-ApbE reactions

The one-pot ApbE–ADPRC reaction was assembled with 0.25 mM $NAD^+$, 0.1 μM ADPRC, 100 μM MBP or NqrC, 10 μM ApbE, 10 mM $MgCl_2$, 20 mM Tris-HCl (pH 9.0), 1 mM EDTA, 100 mM KCl, and a base-exchange substrate. The reaction was incubated at 30 °C for 3 h and subsequently quenched by adding EDTA to a final concentration of 10 mM.

##### 3.7. Enzymatic synthesis and purification of 3-azidopropyl ADP-ribose

Azido-functionalized ADP-ribose derivatives were synthesized enzymatically using *A. californica* ADP-ribosyl cyclase (ADPRC). Reactions were conducted in 50 mM HEPES buffer (pH 7.0) containing 5 mM $MgCl_2$ and consisted of 10 mM $NAD^+$, 2 μM ADPRC, and 2 μL of either 4-pentyn-1-ol or 3-azidopropanol. Reaction mixtures were incubated at 37 °C for 1 h and quenched by heat denaturation at 95 °C for 10 min. Following the ADPRC reaction, excess 3-azidopropanol remained in the reaction mixture and could interfere with subsequent click chemistry reactions. Therefore, an extraction step was performed to purify the ADPRC reaction products. Briefly, 400 μL Milli-Q water was added to 100 μL of the reaction sample to obtain a final volume of 500 μL, followed by the addition of an equal volume of ethyl acetate. The mixture was vortexed thoroughly and allowed to stand until complete phase separation occurred, after which the upper organic phase was removed. This extraction procedure was repeated 12 times. The aqueous phase was then dried using a vacuum concentrator and resuspended in 20 μL Milli-Q water.

##### 3.8. Click chemistry reaction of azido ADP ribose with alkynyl-sulfo Cy3

For click chemistry reaction, stock solutions of 25 mM $CuSO_4$ and 100 mM sodium ascorbate were freshly prepared. $CuSO_4$ and THPTA were pre-incubated at room temperature for 30 min. Subsequently, the click reaction was carried out in 50 mM HEPES buffer (pH 7.0) containing 10 mM sulfo-Cy3-alkyne, 1 mM purified Azido ADPR, 0.2 mM $CuSO_4$ (from the pre-incubated mixture), 1.25 mM THPTA, 5 mM aminoguanidine, and 5 mM sodium ascorbate. Reactions were incubated at room temperature for 1 h. The reaction products were purified using either a C18 gravity column or an ÄKTA chromatography system, followed by concentration through freeze-drying.

##### 3.9. ApbE reaction with sulfo-Cy3 ADP-ribose

ApbE reactions with sulfo-Cy3 ADP ribose were performed with NqrC as a target protein. The reactions contain 1 mg/ml ApbE, 3 mg/ml NqrC, 1 mM sulfo-Cy3 ADP ribose, 1 mM EDTA, 100 mM KCl, and 10 mM $MgCl_2$ in 50 mM Tris-HCl (pH 9.0). Reactions were incubated at 30 °C for 0, 4, or 24 h. Aliquots were collected at the indicated time points, and the reactions were terminated by adding 4× SDS–PAGE loading buffer. Samples were separated on 12% SDS–PAGE gels, and in-gel fluorescence was detected using a GE Typhoon FLA 9500 scanner. Subsequently, the gels were stained with InstantBlue and imaged using a Bio-Rad Gel Doc imaging system.

##### 3.10. Click chemistry reaction of azido ADP ribose with alkynyl agarose

Click chemistry reactions were performed using 200 µL alkynyl agarose suspended in 50 mM HEPES buffer (pH 7.0) in a gravity-flow column. The reaction mixture contained 0.1 mM purified azido-ADPR, 0.2 mM $CuSO_4$, 1.25 mM THPTA, 5 mM aminoguanidine, and 5 mM sodium ascorbate. Reactions were incubated at room temperature for 1 h. Excess azido-ADPR and other small molecules was removed by washing the column with Tris-HCl buffer (pH 9.0).

##### 3.11. ApbE reaction of agarose ADP ribose

To observed ApbE activity towards agarose ADPR, the ApbE reaction was performed with GFP-F2 as target proteins. The reactions was performed in room temperature for 24 h containing 200 µL agarose ADP ribose from click chemistry reactions, 100 µM GFP, 100 µM ApbE, 1 mM EDTA, 100 mM KCl, and 10 mM $MgCl_2$. After the reaction, the columns were washed with 50 mM HEPES buffer (pH 7.0) containing 350 mM imidazole. The immobilized GFP remained bound to the column after washing and was visualized using a Typhoon FLA 9500 imaging system at 473 nm.

##### 3.12. Synthesis of dephospho-CoA-Cy3

Thiol–maleimide conjugation was performed using EZ-Link™ Maleimide-PEG4-DBCO (Thermo Fisher Scientific). Briefly, dephospho-CoA (10 mM) was incubated with Maleimide-PEG4-DBCO (15 mM) in a 1:1 (v/v) mixture of PBS and DMSO for 2 h at room temperature under gentle mixing. The resulting DBCO-functionalized dephospho-CoA intermediate was subsequently conjugated to alkyne-functionalized Cy3 by incubation for an additional 4 h at room temperature.

##### 3.13. ApbE reaction with dephospho-CoA-Cy3

ApbE-mediated labeling of NqrC with dephospho-CoA–Cy3 was carried out in a reaction mixture containing 1 mg/mL ApbE, 3 mg/mL NqrC, 1 mM dephospho-CoA–Cy3, 1 mM EDTA, 100 mM KCl, and 10 mM $MgCl_2$ in 50 mM Tris-HCl buffer (pH 9.0). The reactions were maintained at 30 °C, and aliquots were withdrawn at the indicated time points. Enzymatic activity was quenched by the addition of 4× SDS–PAGE loading buffer. The samples were subsequently analyzed by 12% SDS–PAGE, followed by in-gel fluorescence imaging using a GE Typhoon FLA 9500 scanner. After fluorescence detection, the gels were stained with InstantBlue and documented using a Bio-Rad Gel Doc imaging system.

##### 3.14. Generation of FADylated peptide

FADylated peptides were generated by digesting EUGO with trypsin or LysC. The reactions were carried out using 50 µM EUGO and 0.5 µM trypsin in 50 mM Tris-HCl (pH 8.0), with a total reaction volume of 2 mL, and incubated at 37 °C for 48 h. Digestion efficiency was assessed by SDS-PAGE, and undigested EUGO was removed by heating the reaction mixture at 95 °C for 1 h. The reaction mixture was then dialyzed against water to remove the Tris buffer and subsequently concentrated by lyophilization.

##### 3.15. ApbE reaction with FADylated peptide

ApbE reactions with a FADylated peptide were performed in 50 mM Tris-HCl buffer (pH 9) containing 0.5 mM FADylated peptide, 100 mM KCl, 10 mM $MgCl_2$, 1 mM EDTA, MBP-NF1 (3 mg/mL), and ApbE (1 mg/mL). Reactions were initiated by the addition of $MgCl_2$ and incubated at 30 °C for 24 h. Reactions were terminated by the addition of 4×SDS-PAGE loading buffer. Samples were analyzed by 12% SDS-PAGE, and in-gel fluorescence was detected using a Bio-Rad Gel Doc imaging system. Gels were subsequently stained with InstantBlue and imaged.

##### 3.16. Preparation of ADP-ribosylated F1-actin

Purified G-actin was polymerized by the addition of KCl and $MgCl_2$ to final concentrations of 150 mM and 2 mM, respectively. After 2 hours incubation at room temperature, F-actin was ADP-ribosylated by incubation with recombinant TcART in the presence of $NAD^+$, following the previously described procedure (Belyy et al., 2022). The reaction was performed in 20 mM Tris-HCl pH 8.0, 150 mM NaCl, 1 mM $MgCl_2$ and 1 mM $NAD^+$ for 30 min at 37°C.

##### 3.17. Preparation of ADP-ribosylated G actin

ADP-ribosylation of G-actin was carried out in a reaction mixture containing 47.3 µM G-actin, 1 µM Iota toxin Ia domain, and 100 µM $NAD^+$ in a buffer composed of 20 mM Tris-HCl (pH 7.4), 0.5 mM ATP, 0.2 mM DTT, and 0.2 mM $CaCl_2$. The reaction was incubated at 30 °C for 30 min.

##### 3.18. Preparation of ADP-ribosylated PARP10

The purified GST-PARP10cat was incubated with 150µM NAD+ at 37 ºC for 1 hour to allow automodification, followed by desalting using zeba columns (ThermoFischer 89877).

##### 3.19. Preparation of ADP-ribosylated Ubiquitin

ADP-ribosylated ubiquitin (ADPR-ubiquitin) was generated by incubating 1 µM $SdeA^{519-1100}$ with 25 µM ubiquitin (Ub) and 1 mM $NAD^+$ at 37 ºC for 1 h.

##### 3.20. ApbE reaction with ADP-ribosylated protein

ApbE reactions with ADP-ribosylated F1 actin were carried out in 50 mM Tris-HCl buffer (pH 9) containing 5-10 mg/mL ADP-ribosylated protein (F-and G-actin, PARP10), 10 mM $MgCl_2$, 1 mM EDTA, 100 mM KCl, 3-5 mg/mL target proteins (NqrC, GFP-F2), and ApbE (5-10 mg/mL). Reactions were initiated by adding $MgCl_2$ and incubated at 30 °C for the time intervals indicated in the gel. Reactions were stopped by adding 4× SDS-PAGE loading buffer. Samples were resolved by 12% SDS-PAGE, and in-gel fluorescence was detected using a Bio-Rad Gel Doc imaging system. Gels were stained with InstantBlue and imaged. The reaction involving ADP-ribosylated ubiquitin was performed under the same conditions, with the addition of 250 µM F1 peptide and 5 mg/mL ApbE.
